# Uni-Fold: An Open-Source Platform for Developing Protein Folding Models beyond AlphaFold

**DOI:** 10.1101/2022.08.04.502811

**Authors:** Ziyao Li, Xuyang Liu, Weijie Chen, Fan Shen, Hangrui Bi, Guolin Ke, Linfeng Zhang

## Abstract

Recent breakthroughs on protein structure prediction, namely AlphaFold, have led to unprecedented new possibilities in related areas. However, the lack of training utilities in its current open-source code hinders the community from further developing or adapting the model. Here we present Uni-Fold as a thoroughly open-source platform for developing protein folding models beyond AlphaFold. We reimplemented AlphaFold and AlphaFold-Multimer in the PyTorch framework, and reproduced their from-scratch training processes with equivalent or better accuracy. Based on various optimizations, Uni-Fold achieves about 2.2 times training acceleration compared with AlphaFold under similar hardware configuration. On a benchmark of recently released multimeric protein structures, Uni-Fold outperforms AlphaFold-Multimer by approximately 2% on the TM-Score. Uni-Fold is currently the only open-source repository that supports both training and inference of multimeric protein models. The source code, model parameters, test data, and web server of Uni-Fold are publicly available^3^.

## 1 Introduction

Understanding the three-dimensional (3D) structures of proteins is the preliminary must for studying their functionalities and accordingly the mechanisms of biological activities. Predicting how proteins fold via computational methods has long been a fundamental yet most challenging problem in life science. Along with the development of artificial intelligence, a recent breakthrough on *in silico* protein folding, namely AlphaFold [1], unprecedentedly achieved “near experimental accuracy” on a majority of monomeric proteins. To briefly summarize, this method directly predicts the atomic coordinates of a protein using a combination of its amino acid sequence, multiple sequence alignment (MSA), and solved homologous structures. In AlphaFold, the sequence and MSA information is encoded via Evoformer, an attention-based deep neural network. The predicted structure is decoded via a structure module, which predicts the local frames and torsion angles of all residues.

Adapted from AlphaFold, AlphaFold-Multimer [2] was later developed by the same team via training AlphaFold on multimeric protein structures. AlphaFold-Multimer supports the prediction of protein complex structures with significantly better performances compared with traditional docking methods.

The occurrence of the AlphaFold system undoubtedly shed light on countless new possibilities of life science exploration. Well discussed in [3], these possibilities include assistance in solving experimental structures, structure-based drug discovery, and protein designing, etc. Meanwhile, the system is not yet perfect. For instance, it does not work well in predicting structures of membrane proteins, anti-bodies, and the combinations of proteins and ligands. In addition, a more discouraging fact is that the complexity of the AlphaFold system together with some other inconveniences made it almost impossible for smaller research groups to re-train the system. This expels them from the power of further developing the system, or adapting it to other applications. The aforementioned inconveniences include: 1) the current open-source code of AlphaFold does not contain any training scripts or utilities of the model; 2) the code of AlphaFold is based on JAX framework, which is limited to a community currently much smaller than TensorFlow and PyTorch; and 3) the original AlphaFold was designed and trained on Google Tensor Processing Unit (TPU), which is hardly accessible to the majority of the research community.

In order to encourage wider collaborations in the area, we present Uni-Fold as a thoroughly open-source platform for developing protein folding models beyond AlphaFold. Uni-Fold supports the training and inference of both monomeric and multimeric models with high accuracy and efficiency. In particular, we reimplemented both AlphaFold and AlphaFold-Multimer in the PyTorch framework, and reproduced their from-scratch training processes on larger training data. To summarize, Uni-Fold made the following contributions:

- Uni-Fold is an open-source platform that welcomes community contributions. We proved the correctness of the implementation by reproducing the from-scratch training process of AlphaFold and AlphaFold-Multimer with equivalent or better performances.
- Uni-Fold Multimer is, to the best of our knowledge, the first and only open-source implementation of AlphaFold-Multimer which supports both training and inference.
- With various optimization techniques, Uni-Fold is one of the fastest implementations of AlphaFold. Under similar hardware configurations, the training process enjoys about 2.2 times acceleration compared with the official implementation.

The source code, model parameters, and test data of Uni-Fold are now publicly available. Meanwhile, Uni-Fold as a protein structure prediction service is now available at Hermite, a new-generation drug design platform powered by AI, physics, and computing.

## 2 Method

Besides reimplementing AlphaFold and AlphaFold-Multimer according to the official code, we made several alterations and improvements in Uni-Fold. In the rest of this paper, we refer to the monomeric model as Uni-Fold Monomer, and the multimeric one as Uni-Fold Multimer.

### 2.1 Protein Homology

In this subsection, we describe the process of searching homologous sequences and structures in Uni-Fold. Unless otherwise specified, the same pipeline is used in both training and inference.

#### Genetic Search

We reused the genetic search protocol in AlphaFold and AlphaFold-Multimer. We used JackHMMER [4] with MGnify [5], JackHMMER with UniRef90 [6], and HHBlits [7] with Uniclust30 [8] + BFD for monomers. For multimers, we additionally used JackHMMER with UniProt [9] to search for sequences with species annotations. We used the same hyperparameters of the MSA search tools as AlphaFold. Identical sequences in the MSAs were deduplicated. The MSA block deletion and MSA clustering strategies of AlphaFold were implemented as is.

#### Cross-Chain Genetics

It was widely demonstrated that MSAs with properly paired orthologs encode cross-chain co-evolutionary information of protein heteromers and thus serve as strong indicators of the complex structure. In Uni-Fold Multimer, we adopted MSA pairing, a technique proposed in [10] and later used in AlphaFold-Multimer to build cross-chain genetics. For homomeric chains in a complex, we simply concatenated the duplicated MSAs of each chain; for heteromeric chains, we ranked the MSA rows of each chain by the species similarities to the target sequence, and then concatenated rows of the same rank. Unpaired MSA rows, as well as those with no species annotations, were padded with gap symbols.

#### Template Search

The template search process of Uni-Fold was much like to that of AlphaFold. To be more specific, we used structure templates that were released before April 29th, 2020. In training, the templates were first filtered such that all templates were released before the target. Top *n* = 20 templates (if existed) modeled by the “sum_prob” output of HHSearch were kept and further sub-sampled to *k* = min(Uniform([0, *n*]), 4) templates. In inference, the top 4 templates were used. For multimers, templates were individually searched for each heteromeric chain and sampled together. We did not use cross-chain templates.

### 2.2 Model and Loss Functions

In this subsection, we describe the model and loss functions of Uni-Fold, which were implemented closely following AlphaFold(-Multimer). Alterations are summarized below.

#### Alterations in Model Architecture

We globally replaced the ReLU activation in AlphaFold(-Multimer) to Gaussian Error Linear Units (GELUs) [11], calculated as

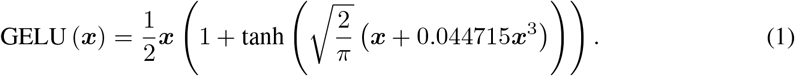

As the `OuterProductMean` module in AlphaFold tended to produce large numerical values which led to training instability, we added a postprocessing layer to its output to lower its values:

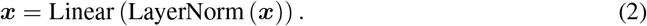

In most auxiliary heads but the predicted-LDDT head, AlphaFold(-Multimer) used a single linear projection (***x*** = Linear (***x***)). We enhanced this with an additional activation function:

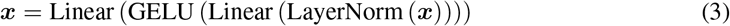

#### Shuffled Multi-Chain Permutation Alignment

In AlphaFold-Multimer, a greedy method was used to disambiguate homologous chains to their respective labels. The method first chose an anchor candidate with the least ambiguity, by which the predicted structure was superposed to the ground truth. A candidate permutation alignment was then derived by greedily minimizing the error of aligned chain centers. The method iterated over all anchor candidates to derive the optimal permutation alignment. In Uni-Fold Multimer, this process was further modified. Instead of greedily minimizing the chain center error, we minimized *C_α_*-RMSD under the superposition of the anchor candidates, so that tangled chains could be better disambiguated. Meanwhile, we shuffled the order of chains before applying the greedy algorithm, and output the best alignment among *n* shuffles.

#### Entity-sharing MSA Mask

Naïvely applying random masks on the MSAs of homomeric sequences in multimers (as AlphaFold-Multimer did) would lead to data leakage in the masked MSA prediction task, as their MSAs were identical. In Uni-Fold Multimer, we addressed this problem with entity-sharing MSA mask, where the same MSA masks were used for chains with identical sequences.

#### Violation Loss

Different violation losses were used between AlphaFold and AlphaFold-Multimer. In both Uni-Fold Monomer and Multimer, we followed the recipe of AlphaFold-Multimer, where the loss of steric clashes of non-bonded atoms was normalized by the number of clashing atom pairs, and the bond angle loss was scaled with weight 0.3.

#### Representation Norm Loss

In both Uni-Fold Monomer and Multimer, we added representation norm losses to encourage numerical stability. The losses 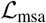 and 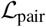 punished the variations of the MSA and pair representations among recycling iterations:

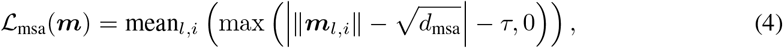

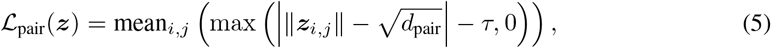

where *d*_msa_ and *d*_pair_ are the dimensions of MSA and pair representations and *τ* = 1 is a tolerance constant.

### 2.3 Implementation and Acceleration

In this subsection, we describe the implementation details of Uni-Fold. Some acceleration techniques are specifically discussed in this subsection.

#### Mixed Precision Training

Mixed precision training [12] is widely used to accelerate the training of large Transformers, where partial or whole calculations of the model is conducted in half-precision to save time and memory. Instead of storing all activations in `bfloat16` format as AlphaFold(-Multimer) did, we used `bfloat16` for most of the layers except for the input embedding layers, geometry-related operations, softmax activations, layer normalizations, and the calculation of all losses. In the specific implementation, parameters in both `bfloat16` and `float32` formats were maintained. After the gradients were calculated, they were copied into `float32` format and then used to update the `float32` parameters. Then, before the next forward process, the `float32` parameters were copied back into `bfloat16` ones. Parameters and gradients were flattened into large tensors during this process, which significantly reduced the time of kernel calls. When casting values from `float32` to `bfloat16`, we used stochastic rounding [13] which was previously shown to encourage numerical robustness. Notably, `float16` was also supported as a feature of Uni-Fold.

#### Operator Fusion

The idea of operator fusion is to merge multiple consecutive operators into one. This accelerates the calculation by reducing the vain cost of repeated global access of GPU memory. Similar to previous works on Transformer acceleration [14, 15, 16], we fused the softmax and layer normalization operators. This operator fusion was particularly important because its two components were done in `float32` format. Fusing them saved not only the memory access time, but also the type converting cost on both time and memory. The operator fusing in Uni-Fold was based on an open-source repository^4^ which was derived from the NVIDIA APEX package^5^. We further optimized its softmax kernel for large columns based on the softmax implementation of OneFlow [17].

#### Memory-Efficient Inference

To reduce the peak memory cost at inference time, AlphaFold proposed a “chunking” technology, which splits the input tensor into multiple small chunks (sub-batches) along one dimension and sequentially forwards the chunks. However, the triangular multiplication module did not use chunking in AlphaFold, as its computation cannot be split in a specific dimension. During inference, Tri-Mul has to allocate 5 times the memory cost of pair representation 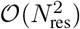, which becomes a memory bottleneck. In order to reduce the burden, we extend the one-dimension “chunking” to the two-dimension “blocking” in Tri-Mul. In particular, we split the computation Tri-Mul into multiple small blocks. For example, with block size 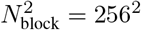, there are ⌈*N*_res_/*N*_block_⌉^2^ blocks. In this way, the peak memory consumption is reduced from 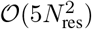 to 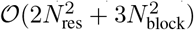. OpenFold[18] uses another solution to reduce the peak memory consumption to 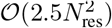.

#### Per-Sample Gradient Clipping

A notable detail in AlphaFold is that the gradient clipping was applied to each sample instead of each batch. However, in most existing AlphaFold replicas such as OpenFold, per-batch gradient clipping was widely used. In Uni-Fold, we implemented both and found that per-sample gradient clipping displays a significant advantage. Detailed comparisons are shown in Section 3.4

#### Distributed Framework and Hardware

We used a cluster of 128 NVIDIA A100 GPUs with 40GB memory for the distributed training of Uni-Fold. The data parallelism strategy of AlphaFold was used, where each GPU contained one training sample at each step. Meanwhile, as mixed-precision and per-sample gradient clipping were used, the distributed algorithm was slightly modified from the standard data parallel algorithm. Specifically, after backward, we copied the `bfloat16` gradients to `float32` ones and then performed per-sample gradient clipping on each GPU independently. An all-reduce operation was finally applied to the clipped gradients.

#### Training Data Compression

In order to reduce run-time parsing costs, we preprocessed and stored sequence features (MSA, templates, etc.) and labels (coordinates from PDB and MMCIF files) as NumPy arrays. To reduce the storage and I/O costs, we adopted several data compression tricks. The PDB dataset consisted of 600,000+ protein chains with only 130,000+ unique sequences. Sharing MSAs and templates among identical sequences reduced the storage space to approximately 1/5. We also compressed the deletion matrices of MSAs into sparse matrices. This further reduced the storage space to approximately 1/6. The features were then compressed to `GZIP` format, which further reduced the storage space to 1/5. A combination of these tricks reduced the storage space from more than 300TB to approximately 2TB with negligible I/O expenses.

## 3 Experiments

### 3.1 Training Uni-Fold

#### Training Protocol

For the training of Uni-Fold Monomer, we used a much simpler two-stage scheme compared with the official AlphaFold. In the initial training stage, we followed the same setting as AlphaFold. In the finetuning stage, we skipped the first finetuning stage of AlphaFold and directly finetuned the model following the configurations of model 1.1.2. Uni-Fold Multimer adopted a similar two-stage scheme, where the initial training and the finetuning configurations of AlphaFold-Multimer were used. Details of the training protocol are summarized in Table 1, where we *italicize* our alterations to the configurations of AlphaFold(-Multimer).

**Table 1:** Training protocol and time for Uni-Fold. Alterations are *italicized*.

| Task Model | Monomer |  | Multimer |  |
| --- | --- | --- | --- | --- |
|  | Init. Training | Finetuning | Init. Training | Finetuning |
| Parameters initialized from | Random | Init. Training | Random | Init. Training |
| Sequence crop size | 256 | 384 | 384 | 384 |
| Number of sequences (MSA + templ.) | 128 | 512 | 128 | 256 |
| Number of templates | 4 | 4 | 4 | 4 |
| Number of extra MSAs | 1024 | 1024 | 1152 | 1152 |
| Batch size | 128 | 128 | 128 | 128 |
| Peak learning rate | 1e-3 | 5e-4 | 1e-3 | 5e-4 |
| Warm-up steps | 1,000 | 500 | 1,000 | 500 |
| Decay steps | 50,000 | N/A | 50,000 | N/A |
| Decay ratio | 0.95 | N/A | 0.95 | N/A |
| Total training steps | 80,000 | 5,000 | 80,000 | 10,000 |
| Ratio of self-distillation samples | 75% | 50% | 50% | 50% |
| Violation loss weight | 0.0 | 0.02 | 0.02 | 0.5 |

#### Training Data

We collected all PDB structures released before January 16th, 2022, among which chain structures were used to train the monomer model, and assembly structures (including those with one chain), the multimer model. Following AlphaFold, we filtered out the structures with resolutions larger than 9 Å, and those with any single amino acid accounting for more than 80% of the sequence. For multimer training, besides the monomer filter, we further filtered out assemblies with more than 18 chains to encourage training stability.

#### Self-Distillation

Uni-Fold adopted the self-distillation strategy of AlphaFold. Similarly, the self-distillation dataset was constructed from Uniclust30 (version 2018_08). To balance data quality and computational cost, we first filtered the sequence clusters in the dataset such that all center sequences have lengths between 200 and 1,024. This left approximately 5 million clusters, which were further used to search for MSAs against Uniclust30, the dataset itself, with HHBlits. Default search parameters are used except for the number of iterations *n* = 3. Among the output MSAs, we first removed those with less than 200 sequences, then removed sequences that appeared in at least two other MSAs, yielding a final dataset of about 360,000 sequences. Predicted structures by an early version of Uni-Fold were used as labels, where structures of residues with Predicted LDDT lower than 50% were masked. Both Uni-Fold Monomer and Multimer used this self-distillation dataset. We did not use multimeric self-distillation samples.

### 3.2 Accuracy Benchmarks

#### Data

We evaluated Uni-Fold and other baselines on recently released protein structures in the Protein Data Bank (PDB) [19]. We collected a total of 1,181 PDB structures released between January 17th and July 14th, 2022. For monomer evaluations, we collected the structures of all sequences and kept those with less than 40% template identity^6^. The left sequences were further filtered so that all sequences have resolutions less than 3Å and lengths between 50 and 1024, yielding a total of 301 unique sequences with 876 structures^7^. For multimer evaluations, we collected assemblies with 2 or more chains, among which all chain had less than 40% template identity. The left was further filtered so that all assemblies have a resolution of less than 3.5 Å and the total number of residues between 50 and 1536, yielding a total of 162 assemblies. The PDB-IDs of the test dataset are publicly available^8^. To test the models’ power of predicting structures of entire assemblies, we did not process the multimeric structures into contacted pairs of chains as AlphaFold-Multimer did. The homology search process is described in Section 2.1. All baselines used the same features.

#### Metrics

For monomer evaluations, as multiple ground-true structures 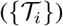 may exist for a sequence, we calculated the metrics using a prediction 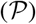 and its best-aligned structure. Taking TM-Score as an example,

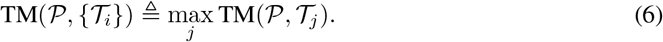

For multimer evaluations, as we evaluated protein assemblies as a whole, using docking-based metrics such as DockQ might lead to confusion. Alternatively, we made a natural extension to adapt single-chain metrics to assemblies by conceptually merging all chains into one. Specifically, for all baselines and metrics on multimeric tasks, the optimal alignment between a prediction and the ground truth was calculated on the entire assembly structure. The scores were then calculated by averaging over all *Cα* atoms. For TM-Score, we used the number of all residues in the assembly to calculate *d*_0_. This process was iterated over all possible permutation alignments and the best score was reported.

#### Baselines

We compared Uni-Fold with AlphaFold(-Multimer) and OpenFold. For monomer evaluations, we report the performances of AlphaFold Model 2 and OpenFold Model 2, which displayed the best accuracy and robustness in early tests. The training of Uni-Fold Monomer also followed the setting of Model 2. For AlphaFold-Multimer, we report all performances of its 5 public models. We used the `v2` parameters of AlphaFold-Multimer.

#### Results

Table 2 shows the results of evaluations. In general, Uni-Fold displays equivalent or better performances compared with AlphaFold and OpenFold on both monomer and multimer tasks. We also evaluated how well multimer models can predict monomer structures by using AlphaFold-Multimer and Uni-Fold Multimer directly on monomer tasks. An obvious drop in performance was observed, which is consistent with the results in [2]. On both monomer and multimer tasks, Uni-Fold Multimer significantly outperformed AlphaFold-Multimer on RMSD and TM-Score. As two models shared the same data pipeline and model implementation, we would conclude that the elevation is obtained using the updated training datasets and the model and loss alterations discussed in Section 2.2. Nevertheless, according to the performances of GDT-HA and *C_α_*-LDDT, predictions from AlphaFold-Multimer may have more accurate localities compared with Uni-Fold.

We present several example predictions from Uni-Fold and AlphaFold-Multimer in Figure 1. Both models have high accuracies on the three examples (TM-Score > 0.90). Predictions from Uni-Fold Multimer have average TM-score 0.984 and *C_α_*-LDDT 0.924, while AlphaFold-Multimer 0.969 and 0.968. The different performances on the two metrics are consistent with the results in Table 2.

**Figure 1:**
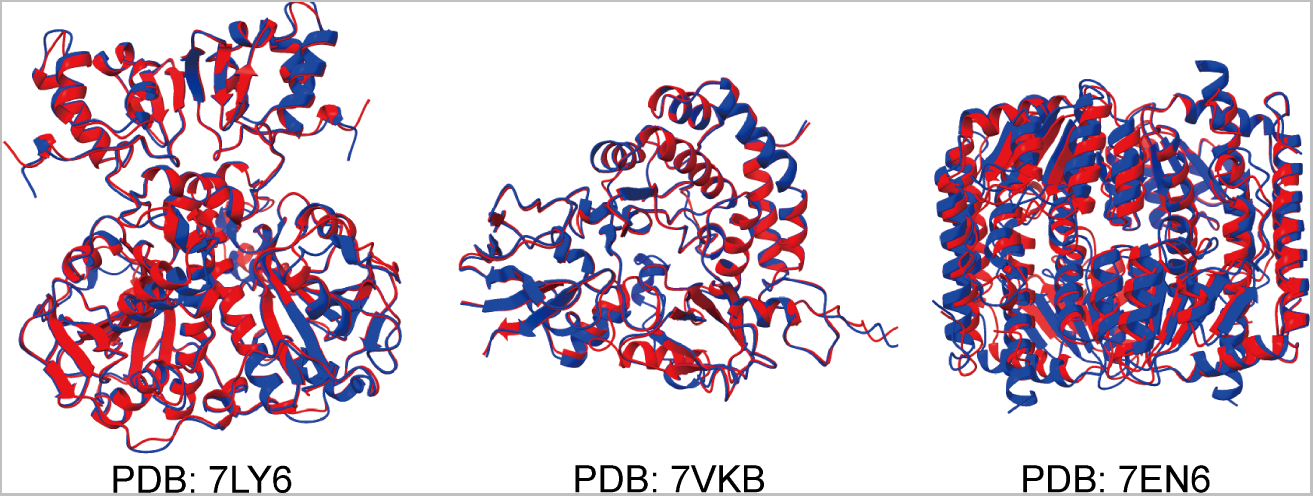
Example predictions from Uni-Fold (blue) and AlphaFold-Multimer (red).

**Table 2:** Accuracy results of Uni-Fold and baselines on recently released PDB structures. *: Note that for multimer models on monomer tasks, the input is given as single sequences instead of assemblies.

| Task | Model | $C_\alpha$-RMSD (Å) | TM-Score | GDT-TS | GDT-HA | $C_\alpha$-LDDT |
| --- | --- | --- | --- | --- | --- | --- |
| Monomer<br>(301 Seqs.) | AlphaFold | **4.669** | **0.776** | 0.686 | 0.531 | 0.879 |
|  | OpenFold | 4.737 | **0.776** | 0.685 | 0.528 | 0.874 |
|  | Uni-Fold Monomer | 4.683 | **0.776** | **0.689** | **0.537** | **0.880** |
|  | AlphaFold-Multimer* | 5.458 | 0.751 | 0.663 | 0.513 | 0.865 |
|  | Uni-Fold Multimer* | 5.142 | 0.756 | 0.658 | 0.496 | 0.864 |
| Multimer<br>(162 PDBs) | AlphaFold-Multimer | 8.263 | 0.763 | 0.607 | 0.458 | 0.883 |
|  |  | 8.115 | 0.764 | 0.605 | 0.456 | 0.882 |
|  |  | 8.086 | 0.764 | 0.606 | 0.456 | 0.883 |
|  |  | 8.141 | 0.764 | 0.611 | **0.462** | **0.885** |
|  |  | 8.238 | 0.761 | 0.609 | 0.461 | 0.882 |
|  | Uni-Fold Multimer | **7.025** | **0.783** | **0.619** | 0.460 | 0.872 |

### 3.3 Efficiency Benchmarks

In this subsection, we introduce the efficiency of Uni-Fold compared with other existing implementations of AlphaFold. On all tasks and all models, mixed precision with `bfloat16` was used. AlphaFold(-Multimer) used 128 TPU cores; MEGA-Fold [20] used 128 Ascend 910; all other baselines used 128 NVIDIA A100 GPUs.

#### Training

We compared the end-to-end training time of Uni-Fold and other AlphaFold implementations. The detailed configurations are displayed in Table 3. On monomer tasks, we followed the model configurations in [21] for fair comparison. On multimer tasks, we followed [2], where *N*_extra_msa_ is inferred from the code. As AlphaFold-Multimer introduced its training details too briefly, we did not know the exact finetuning steps. Results of OpenFold and HelixFold are referred directly from [21], and results of AlphaFold(-Multimer) are from [1, 2]. We do not include the results of FastFold in this table because it did not report training time on 128 GPUs. Table 4 shows the results, in which acceleration ratios to official AlphaFold(-Multimer), “Accel. to AF", are also calculated. Under similar hardware setting and benchmark configuration, Uni-Fold is about 2.2 times faster than the official AlphaFold, also leading other recent baselines. Under the actual configurations of Uni-Fold in Table 1 (Uni-Fold (*real*)), the total training time is approximately 4.11 days. Despite that we do not know the exact steps of AlphaFold-Multimer, we achieved better performances 1.9 time faster.

**Table 3:** Configurations of the total training time benchmark.

| Task | Init./ FT | samples ($\times 10^6$) | $N_{\text{templ}}$ | $N_{\text{res}}$ | $N_{\text{seq}}$ | $N_{\text{extra\_seq}}$ |
| --- | --- | --- | --- | --- | --- | --- |
| Monomer | Initial Training | 10.0 | 4 | 256 | 128 | 1024 |
|  | Finetuning | 1.5 | 4 | 384 | 512 | 5120 |
| Multimer | Initial Training | 10.0 | 4 | 384 | 128 | 1152 |
|  | Finetuning | ? | 4 | 384 | 256 | 1152 |

**Table 4:** Training times (days) of Uni-Fold and other AlphaFold implementations. *: Not rigorous benchmark performance.

| Task | Model | Init. Training | Finetuning | Total time | Accel. to AF |
| --- | --- | --- | --- | --- | --- |
| Monomer | AlphaFold | 7.— | 4.— | 11.— | 1.00 |
|  | MEGA-Fold* | 11.— | 3.5— | 14.5— | 0.76 |
|  | OpenFold | 8.05 | 2.80 | 10.85 | 1.01 |
|  | HelixFold | 5.29 | 2.26 | 7.55 | 1.46 |
|  | Uni-Fold (*benchmark*) | **3.30** | **1.72** | **5.02** | **2.19** |
|  | Uni-Fold (*real*)* | 3.38 | 0.73 | 4.11 | 2.68 |
| Multimer | AlphaFold-Multimer* | 14.— | 2.— | 16.— | 1.00 |
|  | Uni-Fold Multimer* | 7.45 | 1.09 | 8.54 | 1.87 |

#### Inference

Figure 2 shows the inference speed and memory usage of Uni-Fold and other baselines, benchmarked on one Evoformer layer by an NVIDIA A100 GPU with 40GB memory. In all settings, the number of MSAs is set as 512, and `bfloat16` is used. Uni-Fold is consistently faster than all baselines. It also enjoys less peak GPU memory usage, indicating that with the same hardware, Uni-Fold can be used to predict longer sequences. When using the chunk size of 4, the maximum sequence length that an Evoformer layer can accept is about 6720 in Uni-Fold, about 5888 in OpenFold.

**Figure 2:**
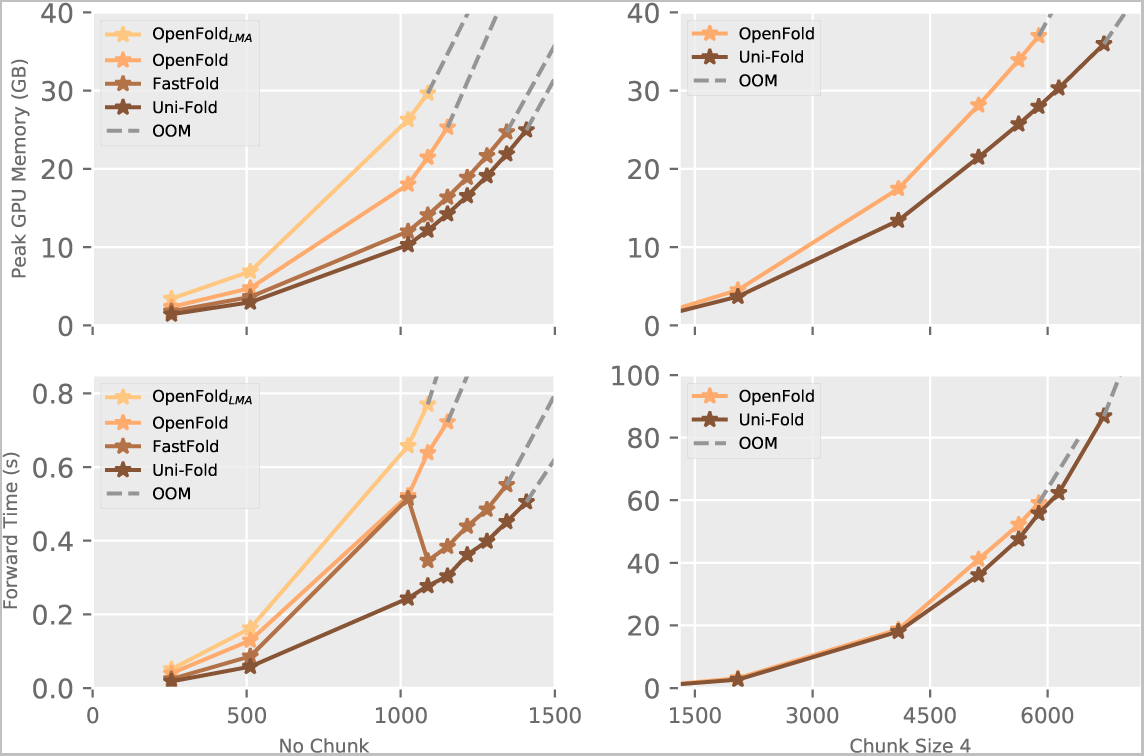
Inference speed and memory usage of Uni-Fold and other baselines with regard to different sequence lengths, on one Evoformer layer. The left is the result without chunking, while the right is the result with chunk size 4. Gray dash lines indicate that the next data points are infeasible due to Out-of-Memory errors.

#### Evoformer Benchmark

To further benchmark the efficiency of Uni-Fold against more baselines that do not include end-to-end training time, following [22], we tested the running time and peak GPU memory consumption of an Evoformer layer in both forward and backward propagation. We used a configuration of 128 MSAs and 256 residues. We included OpenFold and its variant, *OpenFold with Low Memory Attention* (OpenFold_LMA_), as well as FastFold as baselines under this setting^9^. Results are summarized in Table 5, where Uni-Fold displays the best performances on both speed and memory efficiency. A surprising observation is that OpenFold_LMA_ is much slower and uses more memory compared with OpenFold. We double-checked our benchmark scripts and would conclude that the LMA optimization is possibly depreciated in the ongoing development of OpenFold.

**Table 5:** Running time and GPU memory usage of an Evoformer layer. The number of MSA is 128 and the number of residues is 256.

| Model | Forward (ms) | Backward (ms) | Total (ms) | Peak Mem. (GB) |
| --- | --- | --- | --- | --- |
| OpenFold<sub>LMA</sub> | 20.911 | 39.007 | 59.918 | 1.944 |
| OpenFold | 17.198 | 23.703 | 40.901 | 1.836 |
| FastFold | 9.415 | 20.247 | 29.662 | 1.776 |
| Uni-Fold | **8.440** | **18.472** | **26.912** | **1.743** |

### 3.4 Effect of Per-Sample Gradient Clipping

Figure 3 shows the training accuracies measured by *C_α_*-LDDT with regard to different gradient clipping strategies. By default, a per-sample gradient clipping by the global norm with value 0.1 was used. We compared this strategy with per-batch gradient clipping with value 0.1, 0.05, and 0.025. The intuition of reducing the value of per-batch clipping was that per-sample clipping always has smaller gradients than per-batch clipping under the same value. Results in Figure 3 indicate that per-sample clipping leads to consistently better model convergence.

**Figure 3:**
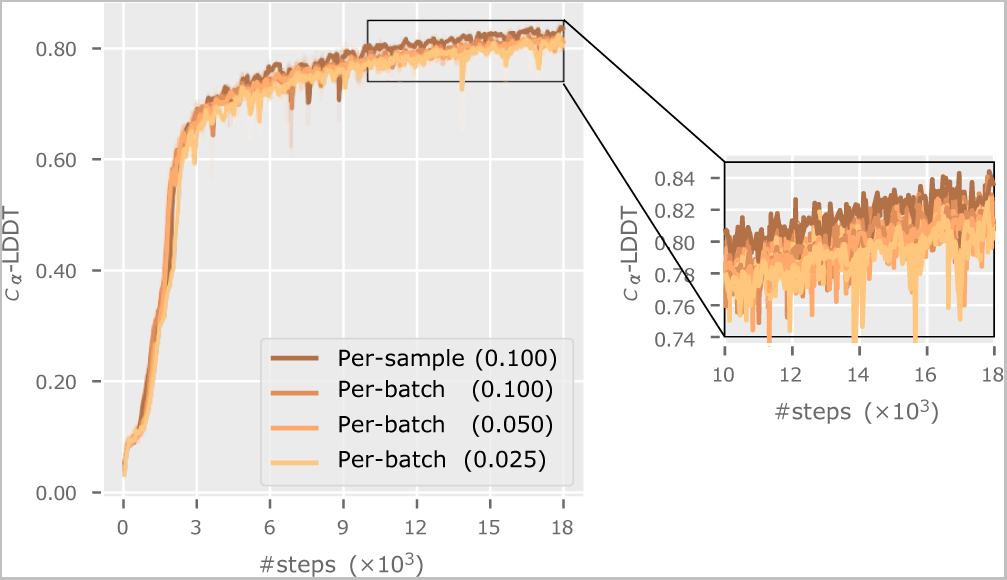
Training accuracies of Uni-Fold Monomer under different gradient clipping strategies.

## 4 Related Work

Plenty of efforts have been devoted to reimplementing or improving AlphaFold. RoseTTAFold [23], known as the earliest re-implementation of AlphaFold (before its release of code), achieved near performance to AlphaFold, while its developers also decided not to release the training code. Open-Fold [18] is an open-source repository that includes the training utilities of the AlphaFold model, yet currently, it does not support the training and prediction of multimeric protein structures. Fast-Fold [22], on the other hand, proposed a model parallelism solution based on the implementation of OpenFold to accelerate training. However, the authors did not provide any details or results of from-scratch training except for efficiency. MEGA-Fold [20] is a recent implementation of AlphaFold trained from-scratch on the MindSpore framework and Ascend 910 hardware. HelixFold [21] is another recent implementation of AlphaFold that supports training and model parallelism under the PaddlePaddle framework.

## 5 The Chronicle of Uni-Fold Development

Released on December 8th, 2021, Uni-Fold v1.0.0^10^ (Uni-Fold-JAX) was the first open-source repository (with training scripts) that reproduced the from-scratch training of AlphaFold with approaching accuracy. Currently, Uni-Fold-JAX is still the only open-source repository that supports the training of official AlphaFold implementation. On April 24th, 2022, we released Uni-Fold v1.1.0 as a service on Hermite. Compared with AlphaFold, Uni-Fold v1.1.0 enjoyed faster training and inference speed as well as better accuracy on newly (at then) released PDB structures. On May 26th, 2022, we released Uni-Fold v2.0.0 on Hermite. Uni-Fold v2.0.0 contained the first reproduction of from-scratch training of AlphaFold-Multimer, with slightly better accuracy on newly (at then) released PDB multimeric structures. On August 1st, 2022, we made Uni-Fold v2.0.0 publicly available on GitHub. The released code supported both training and inference of AlphaFold(-Multimer). Currently, this is the only open-source repository that supports the training of AlphaFold-Multimer.

## Footnotes

3 The source code, model parameters and test data of Uni-Fold are available at https://github.com/dptech-corp/Uni-Fold. The web server is available at the Hermite platform https://hermite.dp.tech. The Colab server is available at https://colab.research.google.com/github/dptech-corp/Uni-Fold/blob/main/notebooks/unifold.ipynb.

4 https://github.com/guolinke/fused_ops

5 https://github.com/NVIDIA/apex

6 The template identity refers to the maximum single template coverage using pipelines in Section 2.1.

7 A sequence may have multiple solved structures in homomers or different assemblies.

8 https://github.com/dptech-corp/Uni-Fold

9 We used OpenFold code with commit ID `a44bbebbfa4dbb8b228e0c8d77338173cf78d699`, FastFold code with commit ID `665e6c97a7d95d3db2df860d104fa3c456c71fe2`.

10 https://github.com/dptech-corp/Uni-Fold-jax

